# Structural basis for cholesterol transport-like activity of the Hedgehog receptor Patched

**DOI:** 10.1101/352443

**Authors:** Yunxiao Zhang, David P. Bulkley, Kelsey J. Roberts, Yao Xin, Daniel E. Asarnow, Ashutosh Sharma, Benjamin R. Myers, Wonhwa Cho, Yifan Cheng, Philip A. Beachy

**Affiliations:** Institute for Stem Cell Biology and Regenerative Medicine, Stanford University School of Medicine, Stanford, California 94305, USA; Howard Hughes Medical Institute, Stanford University School of Medicine, Stanford, California 94158, USA; Department of Biochemistry and Biophysics, University of California, San Francisco, California 94158, USA; Department of Developmental Biology, Stanford University School of Medicine, Stanford, California 94305, USA; Department of Chemistry, University of Illinois at Chicago, Chicago, Illinois 60607, USA; Department of Biochemistry, Stanford University School of Medicine, Stanford, California 94305, USA; Howard Hughes Medical Institute, University of California, San Francisco, California 94158, USA

## Abstract

Hedgehog protein signals mediate tissue patterning and maintenance via binding to and inactivation of their common receptor Patched, a twelve-transmembrane protein that otherwise would suppress activity of the seven-transmembrane protein, Smoothened. Loss of Patched function, the most common cause of basal cell carcinoma, permits unregulated activation of Smoothened and of the Hedgehog pathway. A cryo-EM structure of the Patched protein reveals striking transmembrane domain similarities to prokaryotic RND transporters. The extracellular domain mediates association of Patched monomers in an unusual dimeric architecture that implies curvature in the associated membrane. A central conduit with cholesterol-like contents courses through the extracellular domain and resembles that used by other RND proteins to transport substrates, suggesting Patched activity in cholesterol transport. Patched expression indeed reduces cholesterol activity in the inner leaflet of the plasma membrane, in a manner antagonized by Hedgehog stimulation and with implications for regulation of Smoothened.

---

The Hedgehog (Hh) family of secreted protein signals patterns many tissues and structures during embryogenesis (Chiang et al., 1996; Dessaud et al., 2008; Ingham, 1993) and post-embryonically governs tissue homeostasis and regeneration by regulating stem cell activity (Goodrich et al., 1997; Shin et al., 2011; Teglund and Toftgard, 2010). Impaired Hh signaling is associated with birth defects, including holoprosencephaly (HPE) (Chiang et al., 1996; Roessler et al., 1996), whereas aberrant pathway activation leads to formation of ectodermally-derived cancers such as basal cell carcinoma (BCC) and medulloblastoma (Teglund and Toftgard, 2010). Recent work also reveals that pathway activity in stromal cells actually restrains cancer growth and progression in certain endodermally derived cancers (Lee et al., 2014; Shin et al., 2014).

The mammalian Hedgehog family comprises Sonic hedgehog (SHH), Desert hedgehog (DHH) and Indian hedgehog (IHH), which diverge in their expression patterns but all utilize a common transduction machinery. The major receptor for mammalian Hh signals is Patched1 (PTCH1) (Chen and Struhl, 1996; Fuse et al., 1999; Ingham et al., 1991; Stone et al., 1996), a 12-pass transmembrane protein. PTCH1 suppresses activity of the 7-pass transmembrane protein Smoothened (SMO), thus maintaining Hh pathway quiescence. When PTCH1 is bound by Hh, PTCH1 suppression of SMO is lifted, permitting SMO-mediated pathway activation (Goodrich et al., 1997; Ingham and McMahon, 2001; Ingham et al., 1991) (Fig. S1A). PTCH1 binding also serves to sequester the Hh protein, thus shaping graded tissue responses to Hh signals. Both SMO regulation and Hh sequestration are conserved in metazoans ranging from insects to mammals (Chen and Struhl, 1996; Ingham and McMahon, 2001) and are needed to prevent inappropriate pathway activity. Loss-of-function *PTCH1* mutations account for about 85% of BCC (Johnson et al., 1996), a cancer with over 1 million patients treated annually in the U.S. alone (Rogers et al., 2015), making *PTCH1* perhaps the most commonly mutated tumor suppressor.

Extensive genetic studies have shed little light on PTCH1 biochemical function. Several observations, including homology of PTCH1 to the Resistance-Nodulation-Division (RND) family of bacterial transporters, led to a model suggesting that PTCH1 may act as a transporter that controls access of certain modulatory lipids to SMO, with the binding of Hh acting to inhibit this transport activity (Taipale et al., 2002). Empirical evidence for this model, however, has been lacking. Furthermore, as the PTCH1 sequence is notably different from those of the well-characterized bacterial RND transporters, we do not know whether PTCH1 is truly homologous to RND proteins in its structure and activity.

Structural information regarding mammalian RND homologs is limited. Several distinct structures of another mammalian RND protein, Niemann-Pick type C1 (NPC1), which is required for lysosomal cholesterol efflux, have been reported (Gong et al., 2016; Li et al., 2017; Li et al., 2016b). Extrapolation to PTCH1, however, is complicated by the distinct functions of these proteins and by their considerable divergence in sequence. Whereas PTCH1 comprises 12 transmembrane helices with two major extracellular domains, NPC1 possesses 13 transmembrane helices plus an additional N-terminal extracellular domain essential for its function (Greer et al., 1999). The overall sequence homology in the two topologically related extracellular domains of NPC1 and PTCH1 is low (24.0% and 19.8% amino acid identity for the first and second domains respectively). PTCH1 structural information hence would not only provide mechanistic insights into PTCH1 itself, but might also shed light on the manner in which RND homologs have evolved to execute diverse functions.

We have succeeded in preparing biochemically well-behaved PTCH1 protein that is active in high-affinity Hh protein binding and is suitable for structural investigation. In this study, we report a cryo-EM structure of mouse PTCH1 with an overall resolution of 3.6 Å. This structure reveals a unique dimeric architecture suggestive of positive curvature within the associated membrane. Our analysis of structural elements and topology of the PTCH1 extracellular domains reveals similarities to NPC1 and RND proteins. A particularly striking feature of the PTCH1 structure is the presence of a hydrophobic conduit that courses through the extracellular domain, with cholesterol densities within and at either end of the conduit. These findings together suggest the possibility that PTCH1 may function as a transporter, a suggestion supported by a functional requirement for structural integrity of the conduit, by dramatic changes in inner plasma membrane cholesterol associated with PTCH1 activity, and by rapid ShhN-mediated reversal of these PTCH1-dependent changes.

## Preparation of a stable PTCH1 variant

The full-length mouse PTCH1 protein is poorly expressed and biochemically unstable (Cleveland et al., 2014). We sought to identify a more stable protein variant by using FSEC (<u>f</u>luorescence detection <u>s</u>ize <u>e</u>xclusion <u>c</u>hromatography (Kawate and Gouaux, 2006)) to screen constructs from which potential destabilizing sequences were deleted. Two HECT E3 ubiquitin ligase interaction sites are present in mouse PTCH1, one within the cytoplasmic loop between TM6 and TM7 and the other in the C-terminal cytoplasmic domain (Kim et al., 2015). A previously reported C-terminal truncation, Ptc-CTD (here referred to as Ptch1-A), removes the latter of these HECT E3 sites and boosts PTCH1 expression (Fuse et al., 1999; Kim et al., 2015; Lu et al., 2006). The final construct we selected, hereafter referred to as Ptch1-B, bears an additional deletion that removes the first HECT E3 site within the large cytoplasmic loop and results in an even higher level of expression, and biochemical behavior as a monodisperse peak in FSEC (Fig. S1B).

We measured *in vivo* activity of the Ptch1-B variant by a conventional Gli-dependent luciferase reporter assay in *Ptch1*^−/−^ mouse embryonic fibroblasts (MEF). Due to the absence of *Ptch1* function, these cells show constitutively high activity and no additional activation by Hh stimulation (Taipale et al., 2002). Introduction of the Ptch1-B expression construct suppressed this high basal activity and rendered cells responsive to the ShhN protein ligand, demonstrating that the stabilized Ptch1-B variant maintains *in vivo* activity similar to that of full-length wild-type PTCH1 (Fig. 1A). We then purified Ptch1-B from HEK293 cells using the BacMam expression system (see Methods). Size exclusion chromatography (SEC) of the purified protein revealed an essentially monodisperse near-Gaussian peak, indicating biochemical homogeneity (Fig. S1C). Using microscale thermophoresis, we found that the GFP-tagged ShhN (Sonic Hedgehog signaling domain) binds Ptch1-B at 27±14 nanomolar affinity (Fig. 1B), close to previous measurements of ShhN binding to full-length PTCH1 on the cell surface (Fuse et al., 1999). This result suggests that the purified protein maintains its physiological fold and also indicates that, unlike *Drosophila* Ptc, which requires an Ihog family co-receptor to engage Hh (Zheng et al., 2010), mouse Ptch1 alone is sufficient for high-affinity Hh binding, confirming that the mode of Hh-receptor interaction has diverged between insects and mammals (McLellan et al., 2008).

**Figure 1.**
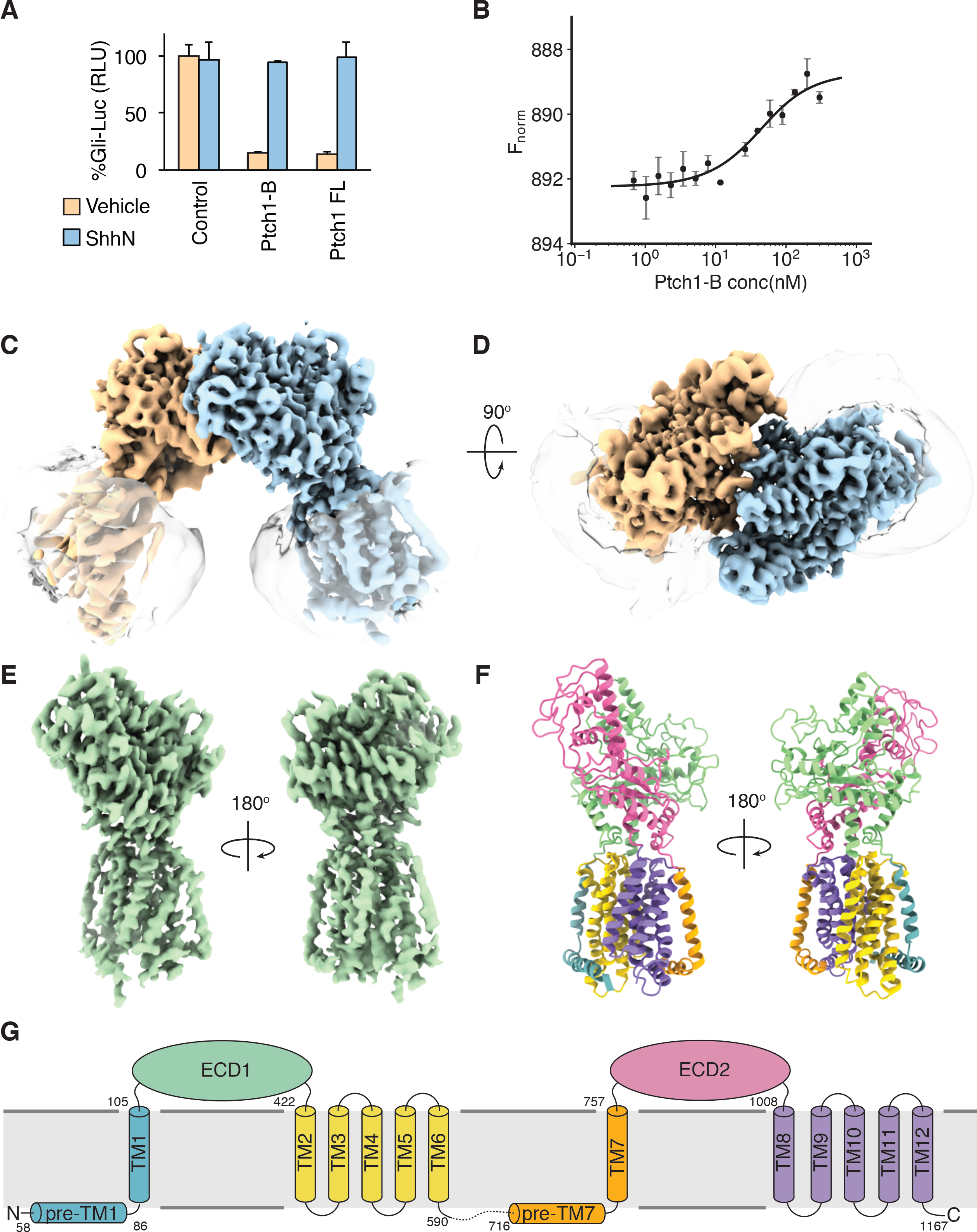
Activity, purification, and structure of the PTCH1 protein. (**A**) Activity of PTCH1 variants was tested in a Gli-dependent luciferase assay in *Ptch1*^−/−^ cell lines. Full length Ptch1 (FL) or the stabilized Ptch1-B variant suppressed high basal activity of *Ptch1*^−/−^ cells and responded to Sonic hedgehog ligand, whereas control cells showed maximal pathway activation. Data for each condition were averaged from triplicates, with error bars indicating standard deviation. (**B**) Affinity of Ptch1-B variant for GFP-tagged ShhN protein was measured in MST. The affinity derived from the standard Kd equation is 27±14 nM. Data for each concentration were averaged from duplicates, with error bars indicating standard deviation. (**C, D**) The electron density map of Ptch1-B sample is shown in side view (**C**, from within the plane of the membrane) and top view (**D**, extracellular perspective), with subunits differently colored. Note the detergent micelles, shown as transparent shells shrouding the transmembrane domains. (**E**) The refined 3.6Å resolution PTCH1 monomer density. (**F)** The PTCH1 atomic model. Domains are colored as follows: pre-TM1 helix and TM1, malachite; extracellular domain 1 (ECD1), lime; TM2-7, yellow; pre-TM7 and TM7, orange; extracellular domain 2 (ECD2), pink; TM8-12, lavender. (**G**) Schematic view of PTCH1 domains, colored as in (**F**).

## Overview of PTCH1 structure

We determined a single particle cryo-EM structure of Ptch1-B (Fig. S2). This structure has an overall resolution of 3.7 Å (Fig. 1C, D and Fig. S3F), and reveals a homodimer with the transmembrane domain of one monomer resolved significantly better than that of the other. The extracellular domains in this map, however, appear symmetrical. Imposition of C2 symmetry improved the overall resolution to 3.5 Å (Fig. S3H), but worsened features in the TM domain of the better-resolved subunit (Fig. S3E, G). Separating each dimer into two individual particles and refining them in concert produced the best density map of the monomer (3.6 Å; map shown in Fig. 1E; resolution in Fig. S3J); this improvement resulted from reducing the uncertainty of monomer positions within the dimer (see below).

Our density maps permitted reliable *de novo* construction of an atomic model of PTCH1 (Fig. 1F and Fig. S4A, B; also see Table 1). In the final model, secondary structures match primary sequence predictions. Furthermore, the positions of 6 cysteines in the extracellular loop between TM1 and TM2 (hereafter ECD1) permit formation of 3 disulfide bonds (density shown in Fig. S4A; schematically drawn in Fig. S4C, D) suggesting that the model correctly threads the primary sequence through the density map, with side chains in appropriate register.

**Table 1.**
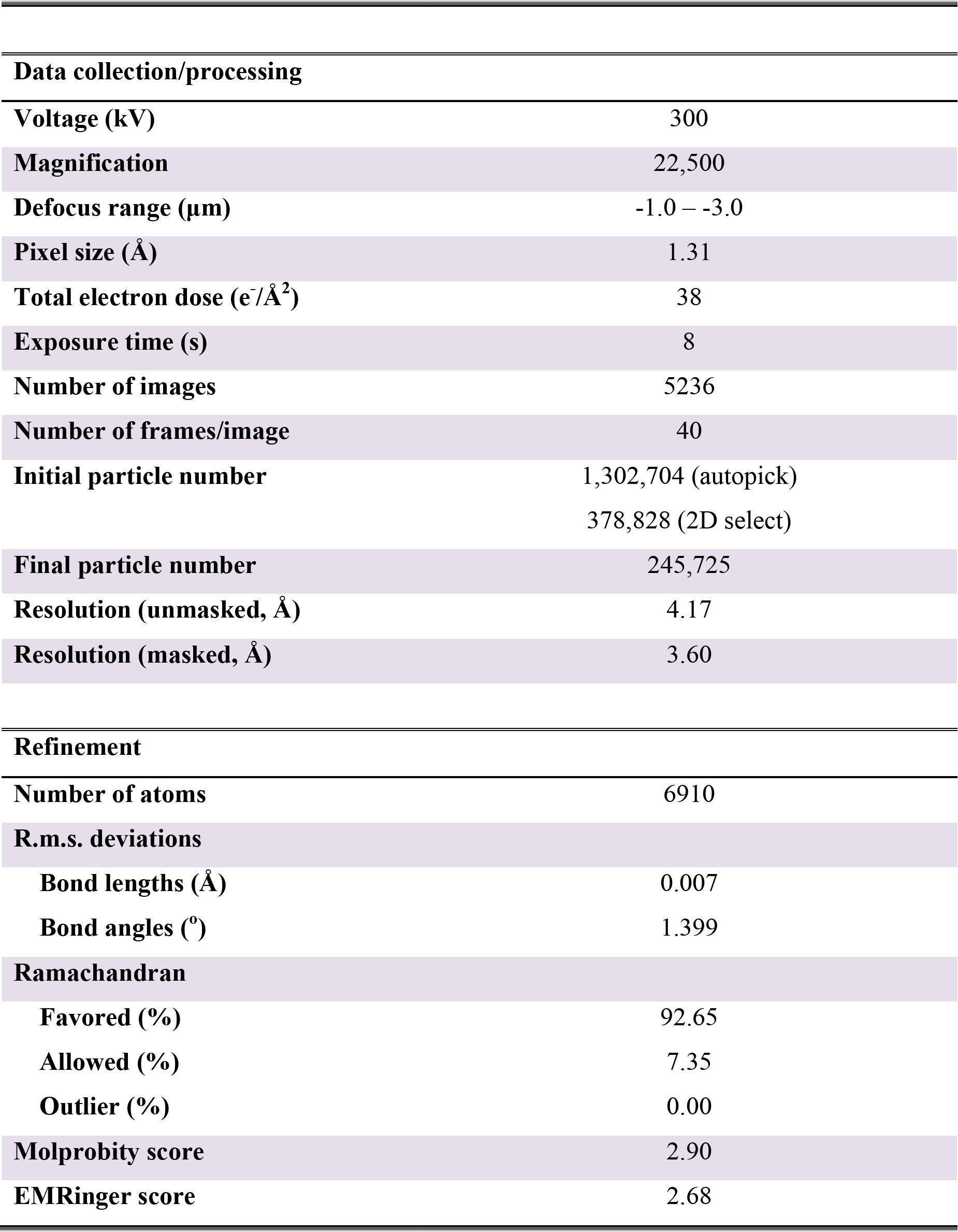
Summary of cryo-EM data collection and model refinement.

Our PTCH1 model exhibits a typical RND transporter-like domain organization (Fig. 1G). Twelve transmembrane helices cluster together to form the transmembrane domain, with pseudo two-fold symmetry between TM1-6 and TM7-12 (Fig. S4E). Two large extracellular domains are located between TM1 and TM2 (ECD1) and between TM7 and TM8 (hereafter ECD2). Most of the intracellular sequence is unresolved, except for two transverse helices preceding TM1 and TM7 at the cytoplasmic face of the protein (Fig. 1F). Although the Ptch1-B protein sample was prepared by ShhN-mediated elution from a ShhN affinity matrix, no density corresponding to the ShhN protein is present in the map, suggesting that the complex of ShhN with the Ptch1-B: dimer is de-stabilized during cryo-EM grid preparation.

## PTCH1 dimer interface

A remarkable feature of our structure is its unique dimeric architecture, in which monomer association is mediated exclusively by the extracellular domains, with the two transmembrane domains projecting away from this vertex at an angle of roughly 50°. This architecture is unusual among membrane proteins of known structure. The angle between the transmembrane domains implies a positive curvature with a radius of ~9 nm in the associated membrane. The dimeric oligomerization state is further supported by SEC-MALS analysis (<u>m</u>ulti-<u>a</u>ngle <u>l</u>ight <u>s</u>cattering) of purified Ptch1-B in amphipol A8-35, which yields a molecular mass of 286 kDa, consistent with a PTCH1 dimer (Fig. S5A).

We constructed a model of the dimer by placing the monomeric atomic structure into the C2 symmetrized density map. In this model, the dimer interface consists of mostly hydrophobic interactions, the tightest being those mediated by residues Y233, I234 and I235 near the center of C2 symmetry (Fig. 2A, B). Other less central residues, however, also contribute (Fig. 2C, D). This combination of tight and loose interactions may preserve the integrity of the dimer while imparting some flexibility at the interface, which may in part account for the lower resolution of one subunit in the non-symmetrized density map. To better reveal the flexibility around the dimer interface, we performed multi-body refinement with a procedure that treats the two subunits within each dimer as rigid body objects to be independently aligned (Fig. 2E and Fig. S2). Based on this analysis, density of the weaker subunit improves significantly while the stronger subunit improves slightly. No major differences in conformation are apparent within monomers at the resolution of our maps (Fig. S5B). The orientations of both subunits within a dimer changed during multi-body refinement, reflecting motion around the dimer interface. As shown in Fig. 2F, the motion includes rotation around both the x and z axes (also see Movie 1). Principal component scores for subunit orientations derived from the multi-body analysis largely follow a distribution of single-peak Gaussian character, suggesting that the motion around the dimer interface is one continuum and does not reflect discrete conformations (Fig. S5C, D).

**Figure 2.**
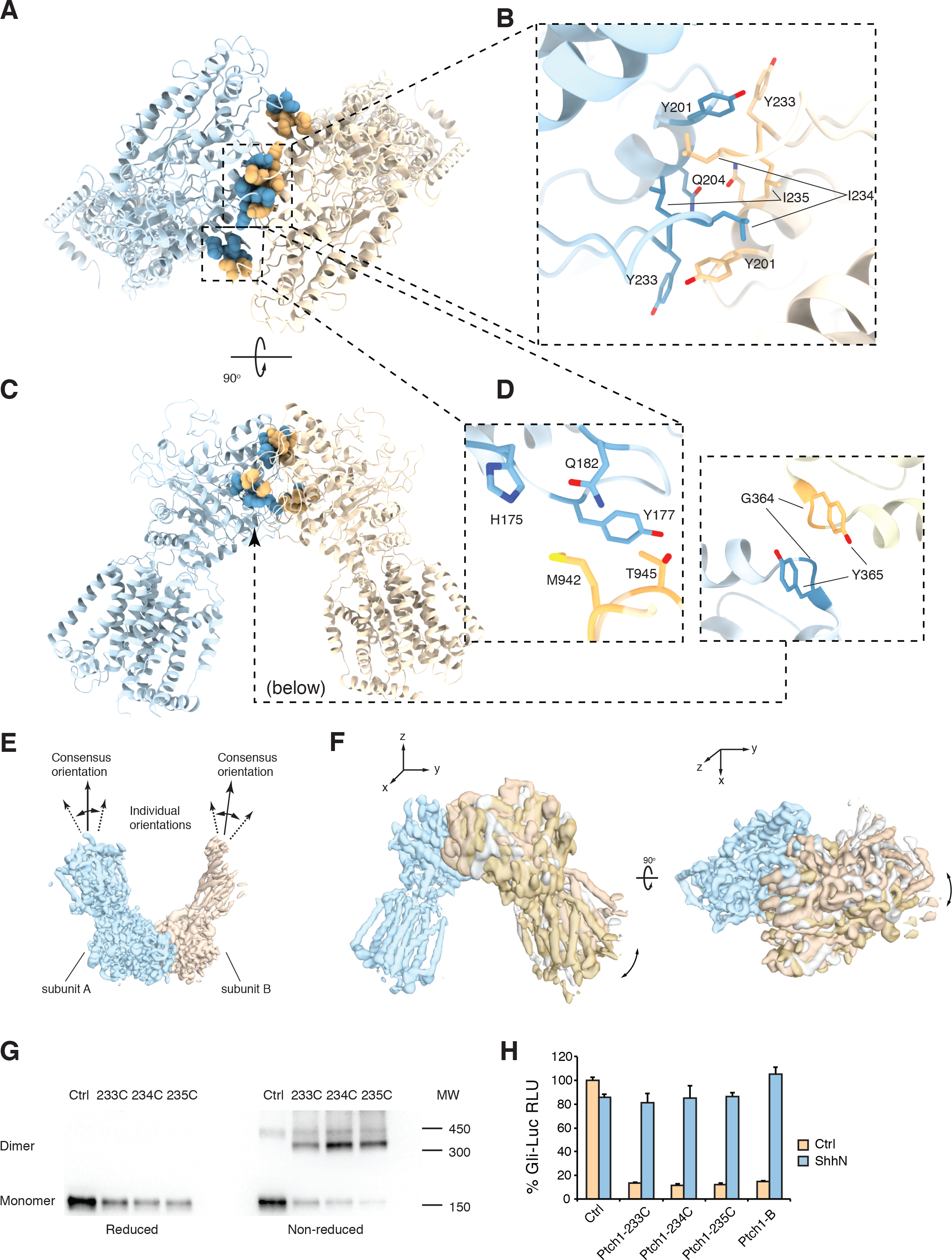
Interactions at the PTCH1 dimer interface. (**A, B**) The PTCH1 dimer interface viewed from the top (**A**) with a close-up view of interacting residues Y233, L234, L235 (**B**). (**C**) Side view of the dimer. (**D**) Enlarged views of residues on the side and the bottom that may stabilize the dimer. (**E**) The two subunits are treated as rigid body objects to be aligned in the multi-body refinement. (**F**) Examples of dimers of distinct relative orientations are superimposed. The blue subunit (left) is aligned as a reference to display relative motion of the other subunit (right). Motions include rotations around both the x and z axes. (**G**) Western blot of HA-tagged Ptch1-B and variants carrying cysteine substitutions at the dimer interface. Under non-reducing conditions, a high molecular weight band corresponding to PTCH1 dimer was seen in all cysteine variants but not Ptch1-B. This band collapsed into a monomer band under reducing conditions. The ~450 kDa band present in all non-reducing gel samples likely is due to non-specific disulfide formation during cell lysis, as 14 cysteines are present on the cytosolic side of Ptch1-B. (**H**) All cysteine-substituted variants retain normal *Ptch1* activity in a Gli-dependent luciferase reporter assay in the *Ptch1*^−/−^ cell line. Data for each condition were averaged from triplicates, with error bars indicating standard deviation.

We tested the *in vivo* occurrence of the PTCH1 dimer by introducing Cys substitutions designed to mediate disulfide bond formation between monomers. Upon cysteine substitution at the residues close to the core of the dimer interface, namely Y233, I234 and I235 (Fig. 2B), we noted by Western blotting the appearance of a band corresponding to a cross-linked dimer, with collapse of this band into a monomer band in the presence of reducing agent (Fig. 2G). This type of reversible cross-linking confirms the proximity of the Cys-substituted residues at the dimer interface. Furthermore, cross-linking of the Y233C variant was less efficient than cross-linking of the I234C or I235C variants, consistent with the relative positions of these residues in the model (the distance between Cα atoms of Y233 in the two subunits is 12 Å, farther than optimal for disulfide bond formation, whereas the distance between I234 or I235 is less than 7 Å, within optimal range for disulfide bond formation). The prevalence of the dimer band in the absence of any oxidizing agents suggests that a significant fraction of PTCH1 exists as a dimer *in vivo*. Because these Cys-subsituted proteins display normal function in SMO suppression and ShhN signal response upon introduction into *Ptch1*^−/−^, MEFs (Fig. 2H), it seems likely that the dimer may remain intact during the normal PTCH1 working cycle. Further work will be required to determine how the membrane curvature implied by this unique dimer architecture may influence PTCH1 localization and function within membrane microdomains.

## Transporter-like features of PTCH1

Comparison of the PTCH1 structure to related structures from other RND homologs reveals essentially the same fold of their transmembrane domains. The PTCH1 transmembrane domain can be superimposed onto those of AcrB, SecDF, HpnN and NPC1 with root mean square deviation (rmsd) values of 6.08 Å, 5.10 Å, 6.85 Å and 1.95 Å, respectively (Fig. 3A and Fig. S6A).

**Figure 3.**
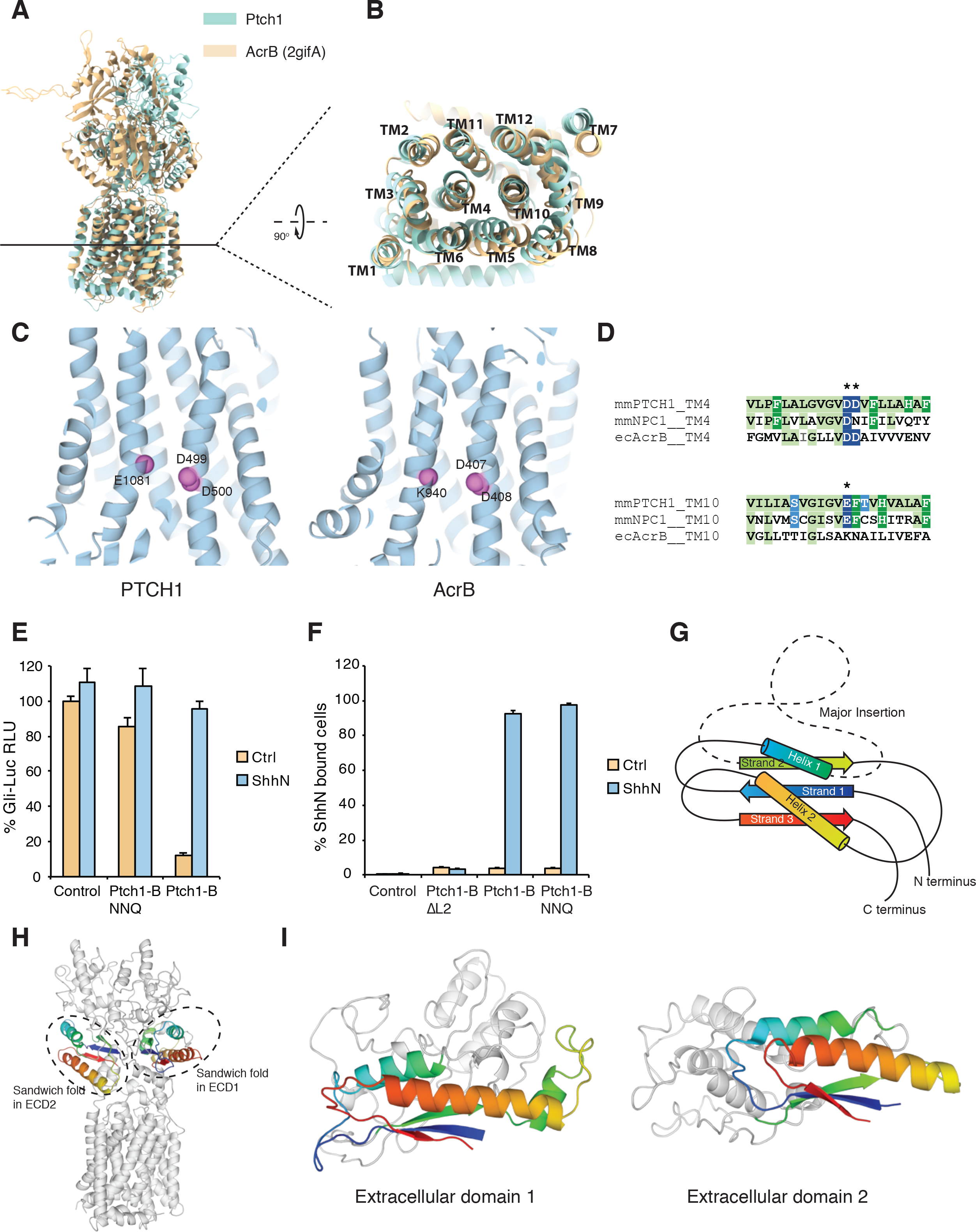
Conserved features of PTCH1 and other RND family members. (**A, B**) Overlay of the PTCH1 structure model (aqua) with AcrB (wheat), shown as seen from the plane of the membrane (**A**) and as a section through the transmembrane domain (**B**). The transmembrane domains align well whereas the extracellular/periplasmic domains diverge significantly. (**C**) The key charged residues in TM4 and TM10 are shown as spheres in the cartoon representations of PTCH1 (left) and AcrB (right) and are highlighted (asterisk) in the sequence alignment of PTCH1, mouse NPC1 and *E. coli* AcrB (**D**). (**E, F**) These charged residues are altered to generate Ptch1-B NNQ (D499N, D500N, E1081Q). This PTCH1 variant failed to suppress basal activity in *Ptch1*^−/−^ cells in a Gli-dependent luciferase assay (**E)**, but still bound ShhN **(F)**. Data for each condition were averaged from triplicates, with error bars indicating standard deviation. (**G**) Schematic view of the conserved alpha-beta sandwich fold present within the extracellular/lumenal/periplasmic domains of RND family proteins. Structural elements are colored in spectral sequence from amino to carboxy-termini (blue to red, respectively), with a common site of non-conserved sequence insertion shown as a dashed line. (**H**) PTCH1 structure highlighting two alpha-beta sandwich folds in the extracellular domain. (**I**) Enlarged view of the sandwich folds in ECD1 (left) and ECD2 (right). The sandwich fold is colored as in (**G**), with inserted sequences shown in gray.

In most well-characterized bacterial RND transporters, the transmembrane domain drives the conformational cycling required for transport, by conducting ion flow through a pathway lined by a triad of charged residues in TM4 and TM10. This charged triad is conserved in PTCH1 (Fig. 3C), and we tested its importance by introducing the charge-neutralizing alterations D499N, D500N, and E1081Q (referred to as “NNQ”). This NNQ variant is no longer able to suppress SMO upon introduction into the *Ptch1*^−/−^, cell line (Fig. 3E; also reported in ref. (Myers et al., 2017)) but retains the ability to bind ShhN (Fig. 3F), consistent with the possibility that the PTCH1 transmembrane domain may utilize ion flow to drive a conformational cycle required for its function.

The extracellular domain of PTCH1 in contrast is more distinctive, containing not only conserved, but also highly divergent features (Fig. S6B). The conserved feature common to the extracellular/periplasmic/lumenal domains of all of these proteins is an “open face” alpha-beta sandwich fold consisting of two alpha helices (the sausages of the sandwich) resting on one face of a beta sheet comprising three strands (the single bread slice of the sandwich). As shown in the topology diagram (Fig. 3G), this conserved feature begins with a beta strand (strand 1), continues as an alpha helix (helix 1), and is then interrupted by insertion of a highly divergent sequence. Following the interruption, the conserved alpha-beta sandwich feature continues with a beta strand (strand 2), a second alpha helix (helix 2), and a final beta strand (strand 3). A variation of this theme is seen in AcrB, in which each periplasmic domain comprises two alpha-beta sandwich domains (Fig. S6B), the first one uninterrupted and the second one interrupted by the “TolC docking domain” (see below). In SecDF, the insertion is between helix 2 and strand 3, rather than between helix 1 and strand 2 (Fig. S6B).

It is noteworthy that amino acid sequences inserted into the conserved alpha-beta sandwich fold are associated with the unique activity of each protein: in AcrB, this insertion (the “TolC docking domain”) docks to the TolC protein conduit for substrate extrusion through the outer membrane (Koronakis et al., 2000; Tamura et al., 2005); in NPC1, the insertion contains the site for binding of NPC2, a docking partner required for cholesterol transfer (Li et al., 2016a), and for binding of glycoproteins of filoviruses including Ebola, which exploit NPC1 for entry into host cells (Wang et al., 2016); in SecDF, the insertion is the head domain, proposed to function in peptide translocation (Tsukazaki and Nureki, 2011). The insertion in ECD1 of PTCH1 mediates monomer association and accounts for the unique dimeric architecture of PTCH1, which is not seen in most bacterial RND transporters. Given the conservation of the transmembrane domain and the presence of the alpha-beta sandwich fold in both extracellular loops, it is likely that the ancestral progenitor of these proteins comprised a six transmembrane protein with a single large loop protruding between TM1 and TM2, and containing one or two sandwich folds; these sandwich folds then evolved further to incorporate various insertions dedicated to particular functions, either before or after duplication of the entire six transmembrane domain to generate the modern twelve transmembrane topology (Fig. 4). NPC1 then acquired an additional transmembrane and luminal domain extension of its amino-terminus.

**Figure 4.**
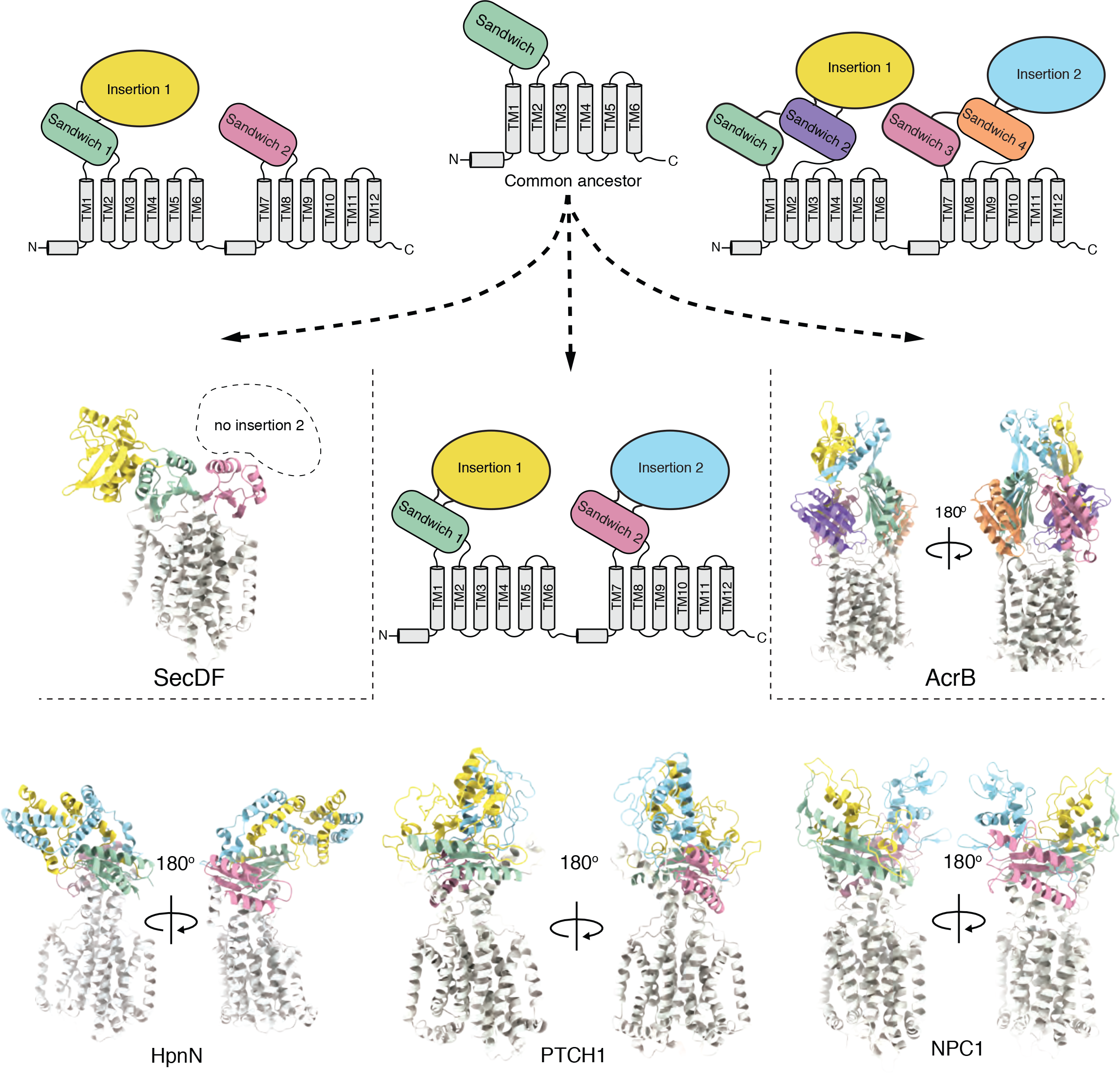
Potential evolutionary derivation of various RND transporters. The ancestral RND progenitor most likely comprised 6 TM helices with one sandwich fold inserted between TM1 and TM2. Duplication of the entire progenitor led to a 12 TM structure. Before or after this duplication, an insertion into the sandwich fold imparted diverse functions, as shown for SecDF, HpnN, NPC1 and PTCH1. In AcrB, duplication of the sandwich fold likely preceded duplication of the 6 TM progenitor, giving rise a total of 4 sandwich folds.

Among bacterial RND homologs of known structure, the overall domain organization of PTCH1 most resembles that of the *Burkholderia multivorans* transporter HpnN, required for hopanoid export to the outer membrane (Kumar et al., 2017). Both PTCH1 and HpnN contain one alpha-beta sandwich fold in each of their extracellular/periplasmic domains, with an interrupting sequence inserted between helix 1 and strand 2 (Fig. 4 and Fig. S6B). In addition, HpnN presents as a dimer during purification and crystallization (Kumar et al., 2017), thus displaying the same oligomeric state as PTCH1 (albeit with a distinct dimer architecture). Dimeric structure thus may not be a particularly rare feature of extended RND family members, even though the most extensively studied member of the family, AcrB, functions as a trimer.

## A hydrophobic conduit within PTCH1 is essential for its activity

To examine the possibility of transporter-like function, we probed for potential substrate-binding cavities in the PTCH1 structure. We found an elongated cavity that courses between the two alpha-beta sandwich folds of the PTCH1 extracellular domain, similar to the path of a cavity in HpnN (Fig. 5A). Interestingly, extra blobs of density are apparent in three locations in high-resolution maps of PTCH1: one near the extracellular end of the cavity (site I), a second within the cavity (site II) and the third one in a groove on the side of TM1 (site III; Fig. 5B). In HpnN, a structurally homologous groove on the side of TM1 has been proposed as the substrate entry site, and is continuous with the cavity in one conformation (Kumar et al., 2017). The residues surrounding these three densities seen in PTCH1 structure are primarily hydrophobic (Fig. 5C), suggesting that the bound molecules are probably hydrophobic, and could be detergent or endogenous lipids carried through purification. The shape and dimension of the densities resemble those of cholesterol (Fig. 5C), and we included cholesterol hemisuccinate (CHS), a commonly used cholesterol analogue, in protein purification (Fig. 5C); the bound molecules thus are likely to be CHS, cholesterol, or its cellular derivatives. These three locations outline a path similar to the substrate translocation path in HpnN, suggesting that a physiological substrate, likely structurally related to cholesterol, may be similarly bound and transported by PTCH1.

**Figure 5.**
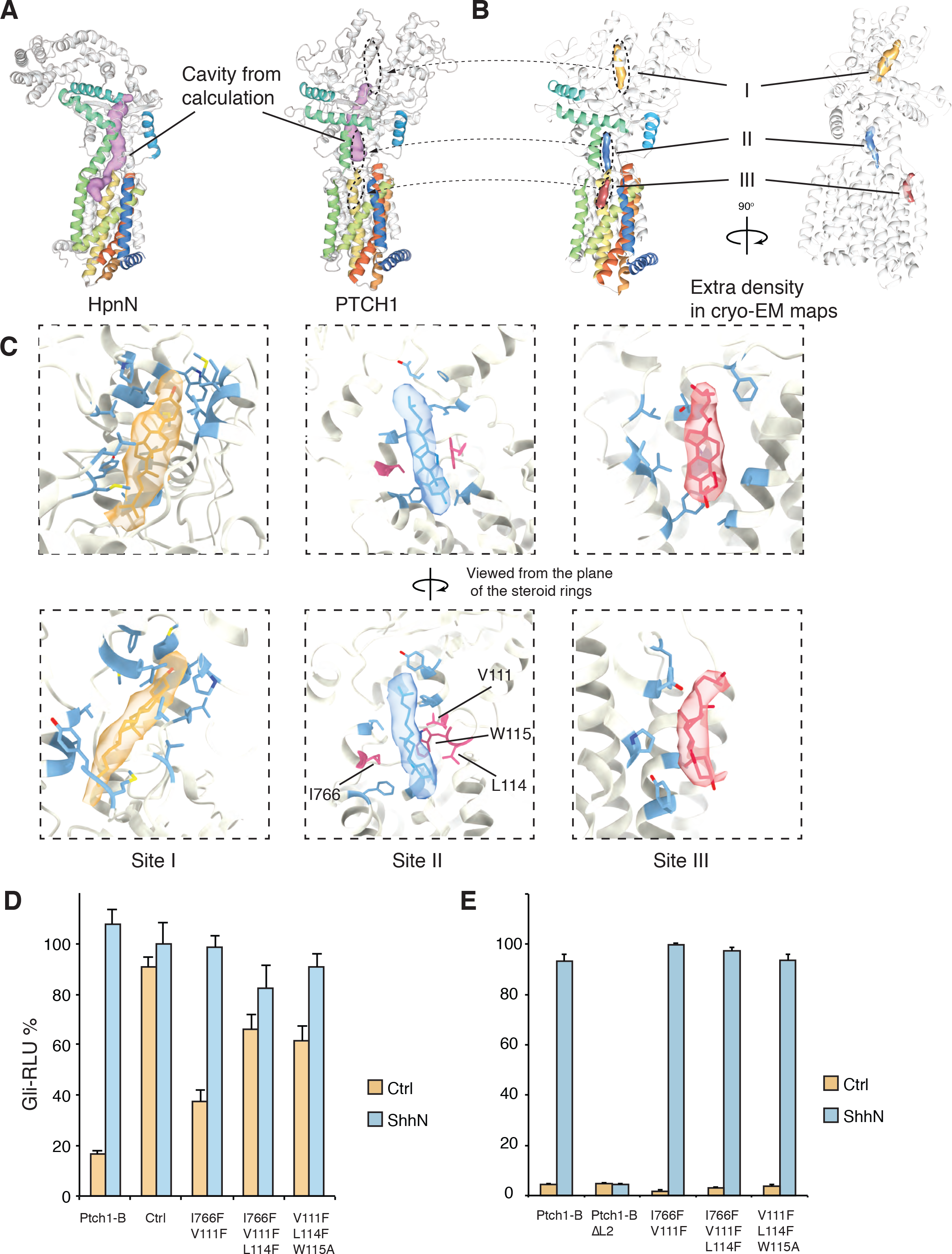
Potential path for lipidic substrate transport in PTCH1. (**A**) Comparison of extracellular/periplasmic domain cavities in PTCH1 and the HpnN hopanoid transporter of *Burkholderia multivorans.* Both cavities are similarly positioned, but the HpnN cavity extends further downward to open to the side of the transmembrane domain. Cavities were calculated using the Caver program. (**B**) Three discrete regions of extra density within (II) and at the ends (I, III) of the PTCH1 cavity are present in high resolution maps. In HpnN, the periplasmic cavity extends to the side of TM1 and is proposed to receive substrates for transport to the outer membrane. Densities I, II, and III outline a potential substrate translocation path similar to that in HpnN. (**C**) Hydrophobic residues surrounding densities I, II, and III favor interactions with hydrophobic substrates. A model of cholesterol is placed inside each density as a reference. (**D, E**) Combinations of residues highlighted in red in (**C**) impaired the ability of PTCH1 to suppress basal activity in Ptch1^−/−^ cells in a Gli-dependent luciferase assay in (**D**), but maintains ShhN binding in (**E**). Data for each condition were averaged from triplicates, with error bars indicating standard deviation.

To test the potential role of this hydrophobic cavity as a conduit for transport, we introduced alterations designed to occlude it. Our efforts focused on site II as this site was included within the elongated cavity and fully enclosed by side chains of hydrophobic residues. We found, in the Gli-dependent luciferase reporter assay in *Ptch1*^−/−^, MEFs described above (Fig. 1A), that combined alteration of hydrophobic residues lining the conduit impaired the ability of PTCH1 to suppress SMO (Fig. 5D) while leaving intact its ability to bind to ShhN (Fig. 5E). Most of these alterations (highlighted in Fig. 5C) substituted bulky phenylalanine residues for aliphatic residues (valine, leucine, and isoleucine), thus potentially blocking the substrate passage by clogging the conduit. This is supported by the retention of ShhN binding, indicating stability and normal folding of these PTCH1 variants. These observations support the hypothesis that a hydrophobic substrate may move through this conduit during the PTCH1 activity cycle.

## Redistribution of membrane cholesterol by PTCH1

As our structure suggests cholesterol transport activity for PTCH1 and cholesterol is known to be required for SMO activation (Cooper et al., 2003), we tested the effect of PTCH1 expression on cellular cholesterol distribution. We have recently developed techniques to directly measure cholesterol activity in the membrane utilizing a set of sensors derived from the cholesterol-binding domain of Perfringolysin O (PFO), each with a distinct affinity for cholesterol and covalently attached to a unique solvatochromic fluorophore (see (Liu et al., 2017) and Methods). The fluorescence emission of these sensors shifts upon interaction with membrane cholesterol (Fig. 6A), thus permitting ratiometric quantification of cholesterol in cellular membranes. The use of a pair of spectrally orthogonal sensors, one microinjected into cells and the other added to the extracellular medium, permits simultaneous *in situ* quantification of cholesterol in the inner leaflet (IPM) and the outer leaflet (OPM) of the plasma membrane. Using these approaches, we recently found that cholesterol in the plasma membrane of mammalian cells is asymmetrically distributed across the bilayer with a level of accessible cholesterol 10-15 fold higher in OPM as compared to IPM in many cell types, (Liu et al., 2017), including HEK293 cells used for our studies here.

**Figure 6.**
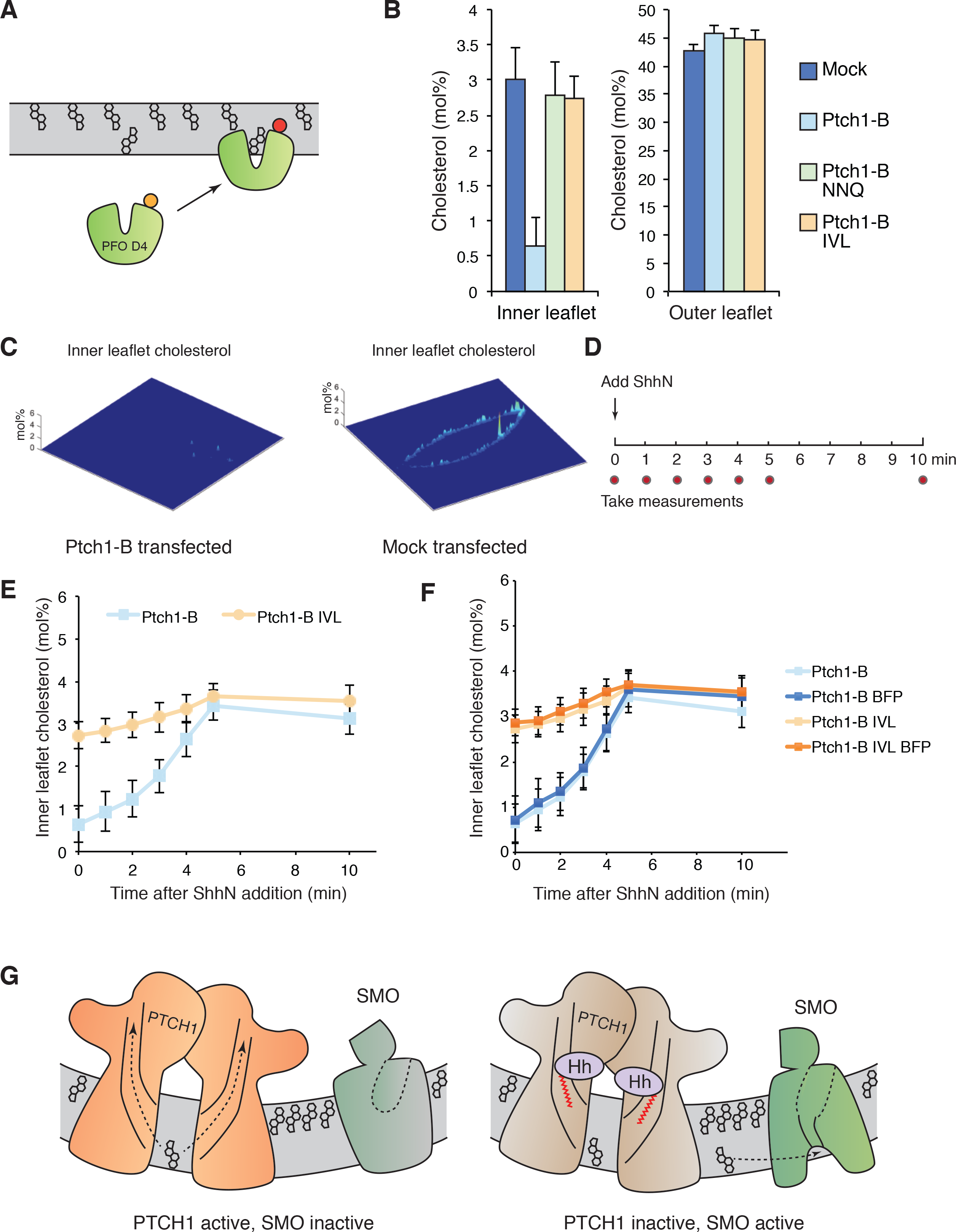
PTCH1 alters cellular cholesterol distribution in the plasma membrane. (**A**) Schematic diagram of the cholesterol sensor. The cholesterol-binding D4 domain from Perfringolysin O was engineered and modified with a solvatochromatic fluorophore such that its fluorescence emission shifts upon binding to cholesterol in the plasma membrane. This engineered sensor provides a ratiometric readout for cholesterol content of the outer leaflet or the inner leaflet (with cell microinjection) of plasma membrane. (**B**) The cholesterol content of the inner (IPM) and outer leaflets (OPM) is compared in the bar graph. The mock transfected cells, or cells transfected with Ptch1-B harboring alterations to the charged triads in the transmembrane domain (NNQ), or blocking mutations in the transport conduit (I766F, V111F, L114F, IVL), showed similar level of inner and outer leaflet cholesterol. In contrast, Ptch1-B transfected cells, the population with lower cholesterol content (approximately half the total cells measured, as in Fig. S7A), showed a dramatic reduction in inner leaflet cholesterol content, whereas the outer leaflet cholesterol content was largely unchanged. For mock transfected cells, Ptch1-B and Ptch1-B NNQ, n=20; for Ptch1-B IVL, n=10. Cholesterol content is derived from the spatially average of the sensor. (**C**) Inner leaflet cholesterol content of representative cells is shown here. Note that in Ptch1-B expressing cells, inner leaflet cholesterol level was reduced to a nearly undetectable level. (**D**) 293 cells expressing Ptch1-B variants were stimulated with ShhN and cholesterol content measured at intervals before and after purified ShhN (C25II) treatment. (**E**) Changes in IPM and OPM cholesterol levels after addition of ShhN. In the subset of Ptch1-B transfected cells with low IPM cholesterol level, ShhN restored IPM cholesterol level to control level within 5 minutes. In cells expressing the Ptch1-B variant harboring blocking mutations in the transport conduit (IVL), IPM cholesterol was within the normal range and didn’t change in response to ShhN. Ptch1-B has a sample size of 20, and Ptch1-B IVL has a sample size of 10. (**F**) BFP tagged Ptch1-B variants were used here, to provide a direct indication of Ptch1-B expression. The change in IPM cholesterol was similar as seen with untagged Ptch1-B. OPM cholesterol was not measured due to spectral overlap between BFP and the extracellular sensor. BFP tagged samples have a size of 10. (**G**) Model for PTCH1-SMO regulation. PTCH1 may transport inner leaflet cholesterol out of the membrane, and thus maintain SMO quiescence. Upon binding to PTCH1, the amino-terminal hydrophobic extension of ShhN occludes the hydrophobic conduit in PTCH1 and blocks PTCH1 transport function. Increased inner leaflet cholesterol then leads to SMO activation. In all panels, error bars indicate standard deviation.

We assayed the effect of PTCH1 on cholesterol distribution in HEK293 cells by transfecting with constructs for expression of Ptch1-B or PTCH1-B variants. As previously reported, the average IPM cholesterol concentration in mock transfected cells was about 3 mol%, as compared to 43 mol% cholesterol in OPM (Fig. 6B). Cells transfected for expression of PTCH1-B exhibited a bimodal distribution of IPM cholesterol, with IPM cholesterol reduced to a nearly undetectable level in approximately half of the cells, whereas OPM cholesterol remained largely unaffected (Fig. 6B, C; entire cell population in Fig. S7A). Cells expressing the PTCH1-B-NNQ variant (an inactive PTCH1 variant, Fig. 3E) or the PTCH1-B-IVL variant with the conduit blocked (I766F, V111F, L114F, the inactive PTCH1 variant shown in Fig. 5D), in contrast, showed little change in IPM cholesterol as compared to mock transfected cells (Fig. 6B). The bimodal distribution in IPM cholesterol in Ptch1-B transfected cells is likely due to the limited efficiency of transfection, as around half of the cells showed high expression of Ptch1-B.

We then found, upon addition of purified ShhN ligand to inactivate PTCH1-B and quantification at 0, 1, 2, 3, 4, 5, and 10 minutes (Fig. 6D), that IPM cholesterol was restored to normal levels within five minutes (Fig. 6E). The increase in IPM cholesterol was synchronized with a comparable reduction in OPM cholesterol (Fig. S7B), suggesting that the cholesterol changes observed upon inactivation of PTCH1 by ShhN might result from cross-bilayer redistribution.

To provide a direct indication of Ptch1-B expression we used blue fluorescent protein (BFP) tagged Ptch1-B variants; with these variants only IPM cholesterol can be measured due to spectral overlap of BFP with the OPM cholesterol sensor. IPM cholesterol levels in BFP-Ptch1-B expressing cells were nearly identical to those of the subpopulation of untagged Ptch1-B transfected cells showing drastically lowered IPM cholesterol concentration (Fig. 6F). Furthermore, the hydrophobic cavity mutant of Ptch1-B (BFP-tagged or untagged) caused neither the reduction of IPM cholesterol in resting cells nor its increase in response to ShhN (Fig. 6E, F), demonstrating the importance of an intact hydrophobic conduit for the effects of Ptch1-B on redistribution of IPM cholesterol. The rapid reversal of the reduced IPM cholesterol upon ShhN-mediated inactivation of PTCH1 suggests that PTCH1 activity likely acts directly to maintain the reduced level of IPM cholesterol.

A model based on these observations would suggest that PTCH1 suppresses SMO activity by reducing IPM cholesterol content through cholesterol transport via the PTCH1 hydrophobic conduit and that Hedgehog binding to PTCH1 inhibits this transport process, allowing IPM cholesterol content to return to normal and permit SMO activation (Fig. 6G). Consistent with this model, recent structural work shows that SMO in its active conformation contains a tunnel opening to the inner leaflet of the membrane (Huang et al., 2018), potentially correlating with increased IPM cholesterol. Furthermore, *in vitro* reconstitution of purified Smoothened protein demonstrates that its activity can be switched on by cholesterol levels in the range of IPM cholesterol measured here upon Patched inactivation (Myers et al., 2017). It is noteworthy that Hedgehog protein signaling activity, but not affinity of receptor binding, depends on a hydrophobic extension from its amino terminus, usually accomplished *in vivo* by palmitoylation (Chamoun et al., 2001; Pepinsky et al., 1998). Interestingly, when this hydrophobic extension is added as an acyl chain *in vitro*, the degree of Hedgehog signaling activity correlates positively with the length of the acyl chain (Pepinsky et al., 1998). We speculate that the amino-terminal hydrophobic extension may be positioned to occlude the hydrophobic conduit when ShhN is bound to PTCH1, with improved signaling ability correlating with longer acyl chain lengths due to improved blockade of the hydrophobic conduit (see Discussion).

## Discussion

The structure of PTCH1 is consistent with the proposed hypothesis that PTCH1 functions as an RND-like transporter to control access of modulatory ligands to SMO (Taipale et al., 2002). The transmembrane domain strikingly resembles that of bacterial RND homologs, and PTCH1 function also requires the charged residue triad within the transmembrane domain, which mediates the electrochemical force required to drive RND transport. These similarities in transmembrane domain structure and key residues required for function among PTCH1, AcrB, and other RND transporters together suggest that PTCH1 may be capable of harnessing energy from ions flowing down an electrochemical gradient to drive conformational switching.

With regard to possible function as a transporter, PTCH1 is unlikely to derive electrochemical force for transport activity from proton gradients, which in mammalian cells are restricted primarily to late endosomes and lysosomes. PTCH1 is localized in the primary cilium (Rohatgi et al., 2007), an organelle with no barrier to ionic communication with the cytosol (DeCaen et al., 2013) and therefore unlikely to accumulate a strong proton gradient. We note however, that certain bacterial RND homologs have evolved to rely on alternative energy sources. For example, the RND transporter that functions in peptide translocation, which in certain prokaryotes is synthesized as a split protein encoded in two genes, SecD and SecF, can be powered by an alternative ion gradient. SecD2/F2, one of the two SecDF protein pairs in the halophilic marine bacterium *Vibrio alginolyticus*, retains proton dependence whereas the second pair, SecD1/F1, has evolved to depend upon a Na^+^ gradient (Ishii et al., 2015). It is especially noteworthy that despite the distinct ion selectivity of these two protein pairs, their sequences are highly similar (70% and 64% similarity for SecD1/2 and SecF1/2, respectively), suggesting that a relatively small change in sequence suffices to alter ion selectivity in RND transporters. The ubiquitous presence of Na^+^ gradients across mammalian cell membranes suggests the possibility of a role for Na^+^ in driving PTCH1 activity, a notion consistent with the Na^+^ requirement for PTCH1 function recently reported in a simplified PTCH1-SMO regulatory system in HEK293 cells (Myers et al., 2017).

Although the PTCH1 extracellular domain differs significantly from that of the well-characterized bacterial transporter AcrB, the cavity positioned between sandwich domains in PTCH1, HpnN, and NPC1 is similar, suggesting a potentially similar transport mechanism. The proposed substrates for HpnN transport are hopanoids (Doughty et al., 2011), bacterial lipids with sterol-like rigid rings. In addition, NPC1 is proposed to be a cholesterol transporter, and is required for efflux of lysosomal cholesterol. The conduit in NPC1 thus may also be utilized for cholesterol transport, but apparently in the opposite direction, in which cholesterol moves from the lumenal domain into the membrane.

In the Hedgehog signaling pathway, cholesterol is required to activate SMO (Cooper et al., 2003), the downstream target of PTCH1. A previous study (Bidet et al., 2011) reported that Patched expression in yeast cells modestly increased extrusion of a fluorescent cholesterol derivative into the external medium. Interpretation of this study, however, is unclear due to use of a bulky fluorophore (BODIPY) substituting for part of the isooctyl side-chain that normally would be inserted into the membrane, and due to the absence of a direct examination of membrane composition. Our direct measurements of endogenous cholesterol in the membrane together with the presence in our PTCH1 structure of cholesterol-like densities within and at either end of a hydrophobic conduit constitute strong evidence that PTCH1 may function in cholesterol transport. Nevertheless, some questions remain. Cholesterol is insoluble in water and would require a sink for export. HpnN delivers lipidic substrates directly to the outer membrane, and NPC1 apparently receives cholesterol from NPC2, a lipid-carrying partner. Because eukaryotic cells lack an outer membrane, PTCH1 export of a cholesterol-like lipid may require a lipid carrier as the sink, suggesting the possible existence of an as yet unidentified lipid carrier protein that partners with PTCH1. In addition, PTCH1 and its downstream target SMO are trafficked into the primary cilia under physiologic conditions (Corbit et al., 2005; Kim et al., 2015; Rohatgi et al., 2007), which effectively limits cholesterol transport activity to the ciliary membrane, a minute fraction of the total cell surface, thus enabling efficient inhibition of SMO with a low level of PTCH1 expression.

Although cholesterol transport is an attractive model, alternative models are also viable. The rapid reversal of reduced IPM cholesterol upon addition of Hedgehog ligand suggests direct action of PTCH1 activity in maintaining reduced IPM cholesterol, but conclusive evidence for PTCH1-mediated cholesterol transport may require *in vitro* reconstitution, which may further require incorporation of an as yet unidentified cholesterol sink. In addition, cholesterol in the plasma membrane is known to be complexed with other proteins and lipids, which could limit interaction with PFO and our engineered cholesterol sensor to a subset of the total cholesterol present (Das et al., 2014). The change in fluorescence thus may arise from a change in availability of cholesterol through indirect effects on a cholesterol-binding partner, in which case PTCH1 may instead act on this partner. Despite these alternatives, the most parsimonious interpretation of our observations is that PTCH1 suppresses SMO activity by directly acting to redistribute membrane cholesterol.

## Acknowledgements

We thank C. Hong, R. Huang and Z. Yu at the HHMI Janelia cryo-EM Facility for help in microscope operation and data collection. We thank A. Müller, W. Kan, and W. Weis for help with the SEC-MALS analysis, and M. Elazar, and J. Glenn for use of equipment. P. Lovelace, S. Weber, and the Stanford Stem Cell Institute FACS Core provided support. This work is partially supported by NIH grants R01GM102498 (to P.A. Beachy) and R01GM098672, S10OD020054, and S10OD021741 (to Y. Cheng), and R35GM122530 to W. Cho. Y. Cheng and P.A. Beachy were investigators of the Howard Hughes Medical Institute.

